## Supporting information for "Irrelevant auditory and tactile signals, but not visual signals, interact with the target onset and modulate saccade latencies"

**S1 Table. Student's t-tests and related tests for all experiments but F.** The statistical analyses to compare the mean median latency $n_{score}$ of each condition with the SOA=0 ms reference condition followed 3 steps: a normality test (Shapiro-Wilk), a variance test (Levene) and a paired Student's t-test. This table reports the outcome of all these tests together with the effect size (Cohen's d).

| Condition | -240ms | -120ms | -60ms | -30ms | +0ms | +30ms | +60ms | +120ms | +240ms | No signal |
|---|---|---|---|---|---|---|---|---|---|---|
| **Experiment A - Beeps & Saccades** | | | | | | | | | | |
| Mean n-score | -1.84 | -2 | -1.65 | -1.49 | -1.32 | -0.41 | 0.65 | 0.38 | 0.01 | 0 |
| Normality (Shapiro-Wilk) | w=0.86; p=0.1179 | w=0.93; p=0.5612 | w=0.93; p=0.5046 | w=0.93; p=0.5230 | w=0.94; p=0.6014 | w=0.93; p=0.4978 | w=0.89; p=0.2352 | w=0.92; p=0.4243 | **w=0.80; p=0.0281** | |
| Variance (Levene) | w=1.62; p=0.2236 | **w=4.76; p=0.0466** | **w=9.69; p=0.0076** | w=3.32; p=0.0900 | | w=0.01; p=0.9257 | w=0.02; p=0.8829 | **w=5.63; p=0.0326** | **w=7.09; p=0.0185** | |
| t-test | t(7)=-1.70; p=0.1336 | t(7)=-2.89; p=0.0235 | t(7)=-2.55; p=0.0380 | t(7)=-1.50; p=0.1783 | | t(7)=13.85; p=0.0000 | t(7)=8.13; p=0.0001 | t(7)=7.88; p=0.0001 | t(7)=7.65; p=0.0001 | t(7)=-6.85; p=0.0002 |
| Effect size (Cohen) | d=1.05 | d=1.51 | d=0.78 | d=0.36 | | d=1.58 | d=3.64 | d=3.88 | d=3.04 | d=2.42 |
| Significance | | * | * | | | *** | *** | *** | *** | *** |
| **Experiment B - Blocked control** | | | | | | | | | | |
| Mean n-score | | | -1.78 | | -1.47 | | 0.47 | | | 0 |
| Normality (Shapiro-Wilk) | | | w=0.91; p=0.4594 | | **w=0.76; p=0.0241** | | w=0.89; p=0.3120 | | | |
| Variance (Levene) | | | w=0.39; p=0.5452 | | | | w=2.05; p=0.1825 | | | |
| t-test | | | t(5)=-2.32; p=0.0677 | | | | t(5)=11.01; p=0.0001 | | | t(5)=-6.96; p=0.0009 |
| Effect size (Cohen) | | | d=0.53 | | | | d=2.75 | | | d=2.84 |
| Significance | | | | | | | *** | | | *** |
| **Experiment C - Background flashes & Saccades** | | | | | | | | | | |
| Mean n-score | -0.52 | -1.05 | -0.77 | 0.92 | -0.17 | 1.26 | 1.84 | -0.03 | -0.06 | 0 |
| Normality (Shapiro-Wilk) | w=0.97; p=0.8737 | w=0.96; p=0.8076 | w=0.92; p=0.4449 | w=0.88; p=0.2041 | w=0.84; p=0.0816 | w=0.97; p=0.9223 | w=0.87; p=0.1441 | w=0.93; p=0.5137 | w=0.97; p=0.9121 | |
| Variance (Levene) | w=0.00; p=0.9465 | w=0.00; p=0.9993 | w=0.17; p=0.6824 | w=3.71; p=0.0745 | | w=0.86; p=0.3690 | w=2.42; p=0.1424 | w=1.77; p=0.2051 | w=2.83; p=0.1144 | |
| t-test | t(7)=-0.98; p=0.3577 | t(7)=-3.90; p=0.0059 | t(7)=-2.45; p=0.0439 | t(7)=3.00; p=0.0198 | | t(7)=4.51; p=0.0028 | t(7)=6.57; p=0.0003 | t(7)=0.47; p=0.6557 | t(7)=0.38; p=0.7154 | t(7)=-0.68; p=0.5212 |
| Effect size (Cohen) | d=0.51 | d=1.30 | d=0.84 | d=1.04 | | d=2.40 | d=3.60 | d=0.23 | d=0.19 | d=0.24 |
| Significance | | ** | * | * | | *** | *** | | | |
| **Experiment D - Strips flashes & Saccades** | | | | | | | | | | |
| Mean n-score | 0.01 | 0 | 0.25 | 0.47 | -1.7 | 1.43 | 0.97 | 0.18 | -0.06 | 0 |
| Normality (Shapiro-Wilk) | w=0.84; p=0.1943 | w=0.86; p=0.2742 | w=0.89; p=0.3919 | w=0.93; p=0.6082 | w=0.89; p=0.3848 | w=0.86; p=0.2554 | w=0.86; p=0.2758 | w=0.85; p=0.2127 | **w=0.75; p=0.0391** | |
| Variance (Levene) | w=0.79; p=0.4080 | w=0.67; p=0.4431 | w=0.59; p=0.4698 | w=1.97; p=0.2101 | | w=0.04; p=0.8568 | w=0.32; p=0.5901 | w=0.22; p=0.6523 | w=0.15; p=0.7155 | |
| t-test | t(3)=2.01; p=0.1383 | t(3)=2.27; p=0.1079 | t(3)=3.55; p=0.0380 | t(3)=3.87; p=0.0305 | | t(3)=10.71; p=0.0017 | t(3)=4.71; p=0.0182 | t(3)=9.36; p=0.0026 | t(3)=3.14; p=0.0517 | t(3)=-4.57; p=0.0197 |
| Effect size (Cohen) | d=1.70 | d=2.09 | d=2.45 | d=2.21 | | d=4.14 | d=2.99 | d=2.87 | d=2.47 | d=2.28 |
| Significance | | | * | * | | *** | * | *** | | * |
| **Experiment E - Touches & Saccades** | | | | | | | | | | |
| Mean n-score | -2.19 | -1.9 | -1.22 | -0.81 | -0.43 | 0.32 | 0.47 | 0.07 | 0.31 | 0 |
| Normality (Shapiro-Wilk) | w=0.98; p=0.9453 | w=0.91; p=0.3660 | w=0.87; p=0.1666 | w=0.93; p=0.5106 | w=0.92; p=0.4406 | w=0.94; p=0.6330 | w=0.93; p=0.5017 | w=0.85; p=0.1059 | w=0.92; p=0.4519 | |
| Variance (Levene) | w=0.01; p=0.9148 | w=0.00; p=0.9458 | w=0.59; p=0.4540 | w=0.67; p=0.4269 | | w=2.58; p=0.1305 | w=0.01; p=0.9211 | w=0.45; p=0.5120 | w=0.00; p=0.9949 | |
| t-test | t(7)=-8.76; p=0.0001 | t(7)=-10.08; p=0.0000 | t(7)=-5.56; p=0.0009 | t(7)=-2.39; p=0.0483 | | t(7)=3.65; p=0.0082 | t(7)=7.25; p=0.0002 | t(7)=2.54; p=0.0386 | t(7)=3.25; p=0.0141 | t(7)=1.81; p=0.1129 |
| Effect size (Cohen) | d=2.54 | d=2.09 | d=1.05 | d=0.45 | | d=0.80 | d=1.35 | d=0.75 | d=1.04 | d=0.91 |
| Significance | *** | *** | *** | * | | ** | *** | * | * | |

**S2 Table. Student's t-tests and related tests of experiment F.**

| Condition | -120ms | -60ms | +0ms | +60ms | +120ms | No signal |
|---|---|---|---|---|---|---|
| **Experiment F - Touches, Beeps & Saccades** | | | | | | |
| **Tactile** | | | | | | |
| Mean n-score | -1.48 | -1.43 | -0.75 | -0.05 | -0.19 | 0 |
| Normality (Shapiro-Wilk) | w=0.98; p=0.9330 | w=0.85; p=0.1661 | w=0.97; p=0.8839 | w=0.97; p=0.8879 | w=0.95; p=0.7692 | |
| Variance (Levene) | w=0.68; p=0.4279 | w=1.60; p=0.2342 | | w=0.03; p=0.8606 | w=2.69; p=0.1322 | |
| t-test | t(5)=-3.85; p=0.0120 | t(5)=-6.40; p=0.0014 | | t(5)=6.05; p=0.0018 | t(5)=3.51; p=0.0171 | t(5)=4.63; p=0.0057 |
| Effect size (Cohen) | d=1.42 | d=1.41 | | d=1.76 | d=1.86 | d=2.67 |
| Significance | * | *** | | *** | * | *** |
| **Auditory** | | | | | | |
| Mean n-score | -2.19 | -1.77 | -1.14 | 0.48 | 0.21 | 0 |
| Normality (Shapiro-Wilk) | w=0.87; p=0.2183 | w=0.92; p=0.4936 | w=0.96; p=0.8509 | w=0.99; p=0.9744 | w=0.88; p=0.2493 | |
| Variance (Levene) | w=0.00; p=0.9900 | w=1.57; p=0.2381 | | w=2.40; p=0.1521 | w=4.30; p=0.0649 | |
| t-test | t(5)=-7.10; p=0.0009 | t(5)=-2.03; p=0.0987 | | t(5)=7.48; p=0.0007 | t(5)=5.03; p=0.0040 | t(5)=7.96; p=0.0005 |
| Effect size (Cohen) | d=2.74 | d=1.12 | | d=3.22 | d=2.37 | d=4.59 |
| Significance | *** | | | *** | *** | *** |
| **Audio-Tactile** | | | | | | |
| Mean n-score | -1.92 | -2.18 | -1.36 | 0.42 | 0.28 | 0 |
| Normality (Shapiro-Wilk) | w=0.80; p=0.0639 | w=0.87; p=0.2223 | w=0.94; p=0.6925 | w=0.97; p=0.8856 | **w=0.77; p=0.0309** | |
| Variance (Levene) | w=1.25; p=0.2906 | w=0.13; p=0.7233 | | w=0.03; p=0.8763 | w=0.16; p=0.7000 | |
| t-test | t(5)=-1.60; p=0.1710 | t(5)=-3.21; p=0.0237 | | t(5)=10.34; p=0.0001 | t(5)=5.49; p=0.0027 | t(5)=7.46; p=0.0007 |
| Effect size (Cohen) | d=0.79 | d=1.79 | | d=3.70 | d=3.01 | d=4.31 |
| Significance | | * | | *** | *** | *** |

### S3 Table. Student's t-tests and related tests for comparisons between experiments.

| Condition | -240ms | -120ms | -60ms | -30ms | +0ms | +30ms | +60ms | +120ms | +240ms |
|---|---|---|---|---|---|---|---|---|---|
| **Experiment B vs. Experiment A** (baseline: *No beep* ) | | | | | | | | | |
| Mean n-score (B) | | | -1.78 | | -1.47 | | 0.47 | | |
| Mean n-score (A) | | | -1.65 | | -1.32 | | 0.65 | | |
| Variance (Levene) | | | **w=9.93; p=0.0084** | | w=0.04; p=0.8456 | | w=2.11; p=0.1718 | | |
| t-test | | | t(7)=-0.53; p=0.6034 | | t(7)=-0.52; p=0.6102 | | t(7)=-0.48; p=0.6394 | | |
| Effect size (Cohen) | | | d=0.29 | | d=0.28 | | d=0.26 | | |
| Significance | | | | | | | | | |
| **Experiment C vs. Experiment A** (baseline: *No signal* ) | | | | | | | | | |
| Mean n-score (C) | -0.52 | -1.05 | -0.77 | 0.92 | -0.17 | 1.26 | 1.84 | -0.03 | -0.06 |
| Mean n-score (A) | -1.84 | -2 | -1.65 | -1.49 | -1.32 | -0.41 | 0.65 | 0.38 | 0.01 |
| Variance (Levene) | w=1.54; p=0.2356 | **w=4.63; p=0.0493** | **w=9.98; p=0.0070** | **w=10.43; p=0.0061** | w=0.16; p=0.6952 | w=0.53; p=0.4798 | w=2.08; p=0.1713 | w=0.46; p=0.5098 | w=0.68; p=0.4239 |
| t-test | t(7)=4.59; p=0.0004 | t(7)=3.68; p=0.0025 | t(7)=3.15; p=0.0070 | t(7)=5.05; p=0.0002 | t(7)=3.69; p=0.0024 | t(7)=6.22; p=0.0000 | t(7)=5.14; p=0.0002 | t(7)=-2.31; p=0.0364 | t(7)=-0.42; p=0.6839 |
| Effect size (Cohen) | d=2.30 | d=1.84 | d=1.58 | d=2.52 | d=1.85 | d=3.11 | d=2.57 | d=1.16 | d=0.21 |
| Significance | *** | *** | ** | *** | *** | *** | *** | * | |
| **Experiment D vs. Experiment C** (baseline: *No flash* ) | | | | | | | | | |
| Mean n-score (D) | 0.01 | 0 | 0.25 | 0.47 | -1.7 | 1.43 | 0.97 | 0.18 | -0.06 |
| Mean n-score (C) | -0.52 | -1.05 | -0.77 | 0.92 | -0.17 | 1.26 | 1.84 | -0.03 | -0.06 |
| Variance (Levene) | w=1.41; p=0.2629 | w=1.12; p=0.3147 | w=0.24; p=0.6322 | w=0.09; p=0.7728 | w=0.00; p=0.9892 | w=1.40; p=0.2637 | w=4.12; p=0.0699 | w=0.34; p=0.5740 | w=1.27; p=0.2865 |
| t-test | t(7)=0.98; p=0.3493 | t(7)=2.34; p=0.0411 | t(7)=2.15; p=0.0573 | t(7)=-0.58; p=0.5720 | t(7)=-3.52; p=0.0055 | t(7)=0.49; p=0.6372 | t(7)=-2.23; p=0.0497 | t(7)=0.78; p=0.4558 | t(7)=0.02; p=0.9873 |
| Effect size (Cohen) | d=0.60 | d=1.44 | d=1.31 | d=0.36 | d=2.16 | d=0.30 | d=1.37 | d=0.48 | d=0.01 |
| Significance | | * | | | *** | | * | | |
| **Experiment E vs. Experiment A** (baseline: *No signal* ) | | | | | | | | | |
| Mean n-score (E) | -2.19 | -1.9 | -1.22 | -0.81 | -0.43 | 0.32 | 0.47 | 0.07 | 0.31 |
| Mean n-score (A) | -1.84 | -2 | -1.65 | -1.49 | -1.32 | -0.41 | 0.65 | 0.38 | 0.01 |
| Variance (Levene) | w=1.93; p=0.1867 | w=3.53; p=0.0812 | **w=16.22; p=0.0012** | **w=5.38; p=0.0360** | w=0.54; p=0.4731 | w=3.85; p=0.0701 | w=0.52; p=0.4834 | w=1.29; p=0.2751 | **w=5.00; p=0.0421** |
| t-test | t(7)=-1.17; p=0.2603 | t(7)=0.34; p=0.7414 | t(7)=1.46; p=0.1651 | t(7)=1.89; p=0.0795 | t(7)=2.90; p=0.0117 | t(7)=1.59; p=0.1339 | t(7)=-0.61; p=0.5497 | t(7)=-1.24; p=0.2360 | t(7)=1.06; p=0.3052 |
| Effect size (Cohen) | d=0.59 | d=0.17 | d=0.73 | d=0.95 | d=1.45 | d=0.80 | d=0.31 | d=0.62 | d=0.53 |
| Significance | | | | | * | | | | |

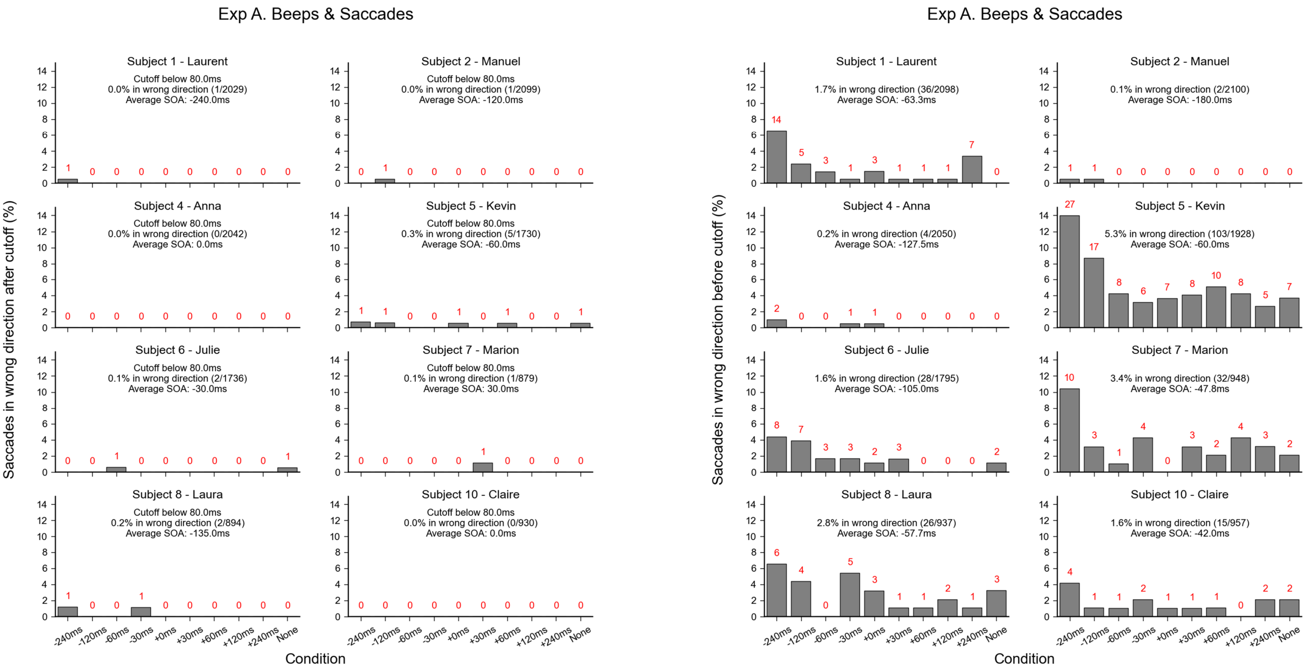


**S4 Fig. Saccades going in the wrong direction of experiment A.** Individual histograms plotting the number of initial saccades going in the opposite direction compared to where the target appeared for each SOA and *No beep* conditions. The **left panel** shows the initial data with all the correctly detected saccades and the **right panel** shows the data after removing those with a gain below 0.4 or with latencies below 80 ms or above 400 ms**.**

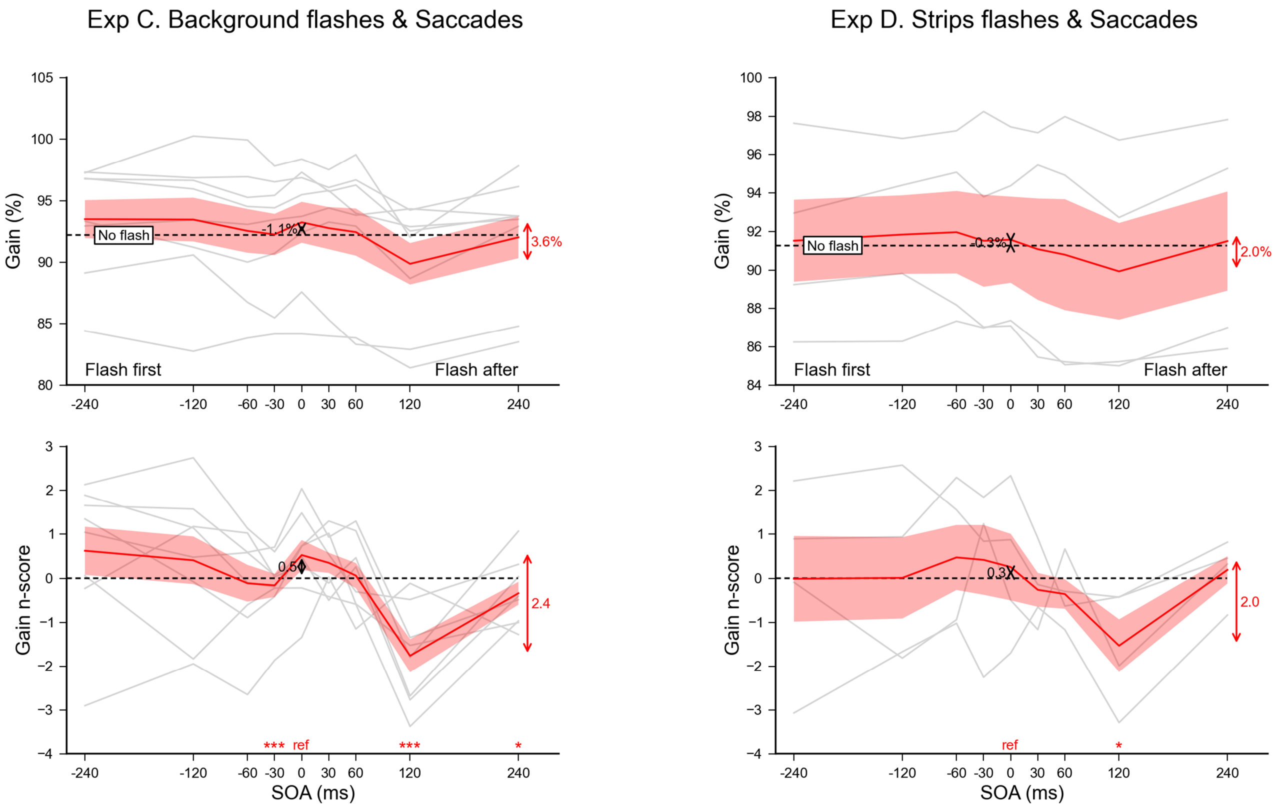


**S5 Fig. Effect of background flashes (left) and strips flashes (right) on saccadic gain.** Saccadic gain averaged across participants for each SOA condition (top) and the corresponding $n_{scores}$ (bottom). Dashed lines show the *No flash* condition level and grey lines show individual results. Error bars indicate inter-individual SEM. Statistics included a single sample t-test performed on the $n_{scores}$ of the SOA=0 ms reference condition to highlight the difference with the *No flash* condition (black arrow), and paired t-tests comparing this reference with each other SOA condition (red stars for each SOA above the X-axis).

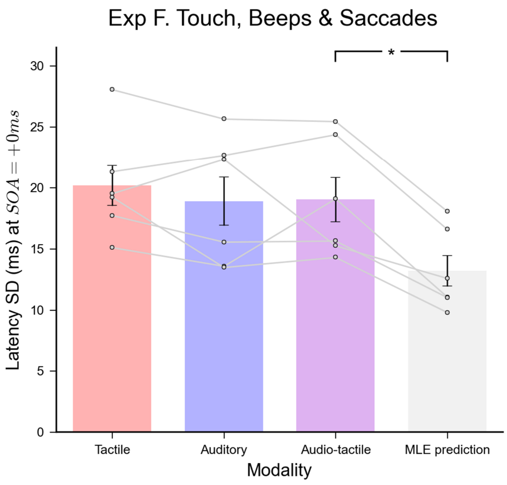


**S6 Fig. Combination of auditory and tactile signals not Bayesian optimal.** Average latency standard deviation computed when SOA=0 ms for each modality with the associated MLE optimal prediction. Comparisons were done using paired t-tests.
